# Diverse processes drive the origination and maturation of super-enhancers/super-silencers during a vast evolutionary timescale of a bicistronic gene

**DOI:** 10.64898/2026.03.09.710545

**Authors:** Nicholas Delihas

**Affiliations:** Department of Microbiology and Immunology, Renaissance School of Medicine; Stony Brook University; Stony Brook, NY 11794 USA

**Keywords:** evolution of super-enhancers, super-silencers, bicistronic gene formation, evolutionary fixation of gene sequence, cultivator gene model, spatiotemporal gene expression, *Alu* TEs in enhancers

## Abstract

A central question in molecular genetics concerns how transcriptional regulatory sequences and *de novo* genes originate and reach evolutionary fixation. In this study, we utilize the human bicistronic gene *SMIM45* as a model to analyze the evolutionary trajectories of gene development. This locus comprises several functional units: three enhancers (one featuring an embedded silencer), an exonic silencer that partially overlaps an ORF, a highly conserved ancestral sequence encoding a 68-aa microprotein, and a human-specific *de novo* gene encoding a 107-aa protein expressed spatiotemporally in embryonic brain tissues. We identify significant disparities in formation mechanisms; for example, the NANOG hESC enhancer originated simply via two *Alu* insertions that constitute the regulatory element. In contrast, the exonic silencer (ATAC-STARR-seq lymphoblastoid silent region 13815), a distinct, novel type of silencer, originated by a combination of diverse mechanisms, including a “cultivator gene” process of base pair fixation consistent with the Cultivator Model proposed by Li Zhao and coworkers. Consequently, *SMIM45* exemplifies novel mechanisms of regulatory element development that occurred over several hundred million years, culminating in the birth of a human-specific *de novo* 107-aa cistron. The properties of these super-enhancers and super-silencers suggest a complex, intricate regulation of the 107-aa protein in fetal tissues.

**Highlights:** *SMIM45*, a bicistronic gene developed an array of enhancers and silencers over several hundred million years of evolution.

Enhancers and silencers formed by diverse processes during very different evolutionary time frames.

The silencer ATAC-STARR-seq lymphoblastoid silent region 13815 in *SMIM45* appears novel, as it is composed of two distinct segments that exhibit disparate origins and developmental pathways.

This intricate regulatory apparatus points to a complex regulation of the 107-aa protein gene expressed in human embryonic brain tissues.

## Introduction

Bicistronic genes are uncommon in eukaryotes, but improved detection techniques are revealing them more frequently [1–3]. These genes form a heterogeneous group with diverse mechanisms for RNA transcription and protein expression. For instance, some utilize alternative RNA transcript isoforms, such as those expressed spatiotemporally in hippocampal neurons [4], while others employ leaky scanning from an internal ribosome entry site (IRES) [5] or an upstream open reading frame (uORF) that inhibits cap-dependent translation [6]. *SMIM45* is also a bicistronic gene, encoding both an ancient 68-aa microprotein and a human-specific, de novo 107-aa protein [7]. Notably, while the 68-aa microprotein is expressed in somatic tissues, the 107-aa protein is expressed spatiotemporally in embryonic brain tissues [8, 9]. A significant feature of *SMIM45* is that it contains of an array of enhancers and silencers, termed super-enhancers and super-silencers that likely regulate the transcription of the 107-aa cistron from its promoter, potentially representing a distinct transcriptional/translational mechanism for a bicistronic gene. This distinguishes the process from protein expression from a bicistronic transcript that relates to the above-mentioned processes.

Enhancers and silencers are short regulatory sequences that are found in great abundance throughout the human genome [10]. They bind transcription factors and can function by regulating cell specific transcription of genes in embryonic tissues [11, 12]. Super-enhancers denote the presence of multiple enhancers in a genomic locus that ensure gene expression in specific tissues. [13–16]. Super-silencers are the presence of two of more silencers in a gene locus that act together to provide strong signals for repression of gene expression whereby repression is dependent on high CpG content of the locus [17, 18].

Because of the array of regulatory elements present in *SMIM45*, the gene lends itself to an analysis of the evolutionary formation of enhancers and silencers. In this paper, we analyze the mechanisms of origination and the timeline of appearance and completion of the super-enhancer and super-silencer regulators during evolution. The study shows regulatory elements formed through diverse mechanisms, highlighting the *SMIM45* gene’s development via the continuous birth of functional elements over ∼400 million years. In particular, our analysis reveals that the short, exonic silencer (a segment of silencer ATAC-STARR-seq lymphoblastoid silent region 13815, which overlaps the C-terminal sequence of the 68-aa protein gene, originated and matured through a combination of distinct molecular processes. This silencer is unusual, as it spans both a gene promoter and an ORF and appears to be unique among known silencing elements. Also presented are comparisons made with known properties of silencers and enhancers that provide insight into how the expression of the 107-aa cistron may be regulated.

Emera *et al* [19] investigated the evolution of enhancers, introducing a model of proto-enhancers as small, early developmental sequences that serve as nucleation sites for further development. We address the role of proto-enhancers/proto-silencers in *SMIM45* development but note that the Emera *et al* model aligns with the previously described evolution of the 107-aa protein cistron. Here, development proceeds by the initial formation of a proto-gene, a short amino acid (aa) sequence called an early developmental sequence [7]. In this framework, the proto-gene originates in ancient species and matures through the contiguous fixation of nearest-neighbor bases of the original as well as secondary proto-genes.

### Background on *SMIM45*

Given the complex nature of the *SMIM45* gene and its regulatory landscape, a comprehensive background is provided. The *SMIM45* ultra-conserved 68-aa microprotein cistron is a pre-existing ancient gene that can be considered a cultivator gene. Lee *et al* [20] presented a model to explain origination of regulatory elements and *de novo* genes based on a proposed cultivator gene, a pre-existing gene that enables the fixation of DNA sequences. On the 3’ side is a *de novo* 107-aa protein cistron found only in humans (Fig. 1). The two *SMIM45* cistrons are linked to and separated by a transcriptional silencer, *LOC130067579,* ATACSTARR-seq lymphoblastoid silent region 13815 [21] whose sequence partly overlaps with the 68-aa C-terminal end. We term this overlapping sequence an exonic silencer. The remaining segment of the silencer, termed here silencer b resides in the intervening sequence, the 107-aa cistron promoter (Fig. 1). Given that enhancer sequences overlapping coding regions are designated as exonic enhancers [22], we term the similarly situated silencer sequence an exonic silencer. About one fourth of all protein genes contain enhancers [22]. An analysis of silencer regions identified in HepG2 cells reveals about 4% of silencers are found in exons [23]. There appears to be no information on silencers that partially overlap open reading fames (ORFs).. The enhancers are designated as enhancers 1-3 and are approximately evenly spaced in the *SMIM45* locus by ∼4kbps (Fig.1). Thus, the *SMIM45* gene and its RNA transcripts have complex arrangements, with transcripts carrying different enhancer sequences in exons or introns (Fig. 1).

**Figure 1.**
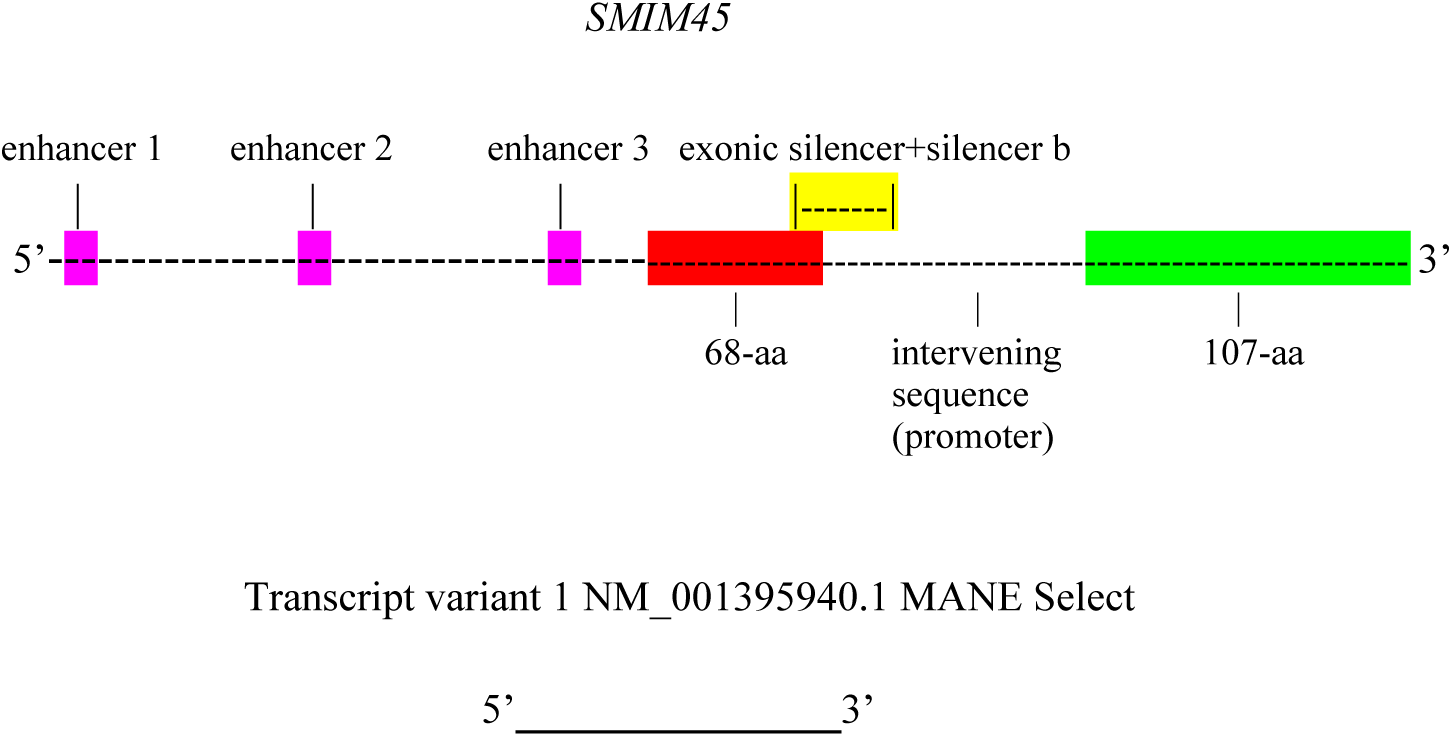
The*SMIM45* gene is 11,991 bps in length and contains three enhancers, the 68-aa protein ORF, the exonic silencer, the 107-aa promoter and the 107-aa protein ORF. Enhancer designations: Enhancer 1, LOC130067578 ATAC-STARR-seq lymphoblastoid active region 19151; Enhancer 2, LOC127896429 H3K4me1 hESC enhancer that also contains an embedded silencer; Enhancer 3, LOC127896430 NANOG hESC enhancer. The exonic silencer is the segment of the transcriptional silencer LOC130067579, ATACSTARR-seq lymphoblastoid silent region 13815 that overlaps the 68aa sequence by 38 bases. Silencer b is the segment of the silencer that overlaps the intervening promoter sequence by 192 bases. The yellow highlighted segment denotes the entire LOC130067579 length., The promoter in the intervening sequence is inferred. Transcript variant 1 is 1540 nt in length and encompasses the *SMIM45* sequence from the 68-aa to the end of the 107-aa sequence. It includes the exonic silencer but no enhancer sequences, however, the intron of processed transcript variant 1 contains the enhancer 3 NANOG hESC. Transcript variant 1 is termed a MANE Select transcript as it annotated in both the NCBI and *Ensembl* data bases with agreement on 5’ end start sites. The 13 other transcript variants also contain the sequence from the 68-aa to the end of the 107-aa sequence and several carry enhancer 1, ATAC-STARR-seq lymphoblastoid active region 19151 in their exons.

The initial study by An et al [24] that described spatiotemporal expression of the *SMIM45* 107-aa protein was partially compromised; the annotation by the *Ensembl* and GenBank data bases for the *SMIM45* locus had not yet been updated from the previous annotation as a lincRNA gene and secondly, detection of a protein was not originally reported. These have now been updated and are no longer issues. With the use of human embryonic cells, the expression of a protein from the 107 aa cistron was shown by multiple experimental techniques. Using human fetal brain cells, Chuan-Yun Li and co-workers [8] reported a protein product from the 107 aa ORF by mass spectrometry and Western blot analysis using specific polyclonal antibody. One of the peptides found, GSGLELVR represents part of the early developmental proto-gene sequence formed during evolution of the 107-aa protein [7]. The immunochemistry analyses show expression in human cerebral cortex and human cortical organoids grown from human embryonic stem cells. In addition, translation from the 107-aa ORF was detected by analysis of public Ribo-Seq datasets where ribosome footprint signals were found from cortical organoids grown from human embryonic stem cells [9]. In other independent gene expression studies, analysis of total RNA from various human tissues reveals RNA expression from *SMIM45* in fetal brain tissues (NCBI BioProject: <u>PRJNA280600</u>) [https://www.ncbi.nlm.nih.gov/gene/339674]. However, a 107-aa cistron transcript has not yet been isolated.

Table 1 lists enhancers found in the *SMIM45* locus together with several known properties of related enhancers. All three show functions that can pertain to regulation of expression of the 107-aa protein. For example, Enhancer 2 related enhancers can act as both enhancers and silencers of gene expression, which matches with the spatiotemporal expression of the 107-aa protein. Translation of the 107-aa protein was detected in human embryonic cerebral cortex and human cortical organoids [8]. Embryonic cerebral cortex tissues consist of proliferating stem/progenitor cells [25]. Enhancer 3 related enhancers bind transcription factor NANOG; NANOG is known to activate several gene promoters in embryonic stem cells [26].

**Table1.** Properties of enhancers related to enhancers 1-3.

| Enhancer | General Properties | Reference |
| --- | --- | --- |
| Enhancer 1<br>*LOC130067578* ATAC-STARR-seq lymphoblastoid active region 19151 | Binds transcription factor activators and silencers within chromatin-accessible regions | [21] |
| Enhancer 2<br>*LOC127896429* H3K4me1 hESC enhancer | Active in embryonic stem cells and associated with activation of gene expression | [27] |
| Enhancer 2<br>*LOC127896429* embedded silencer, silent region_13814 | Binds repressor transcription factor s and prevents transcription; essential to secure gene expression in specific tissues at specific times (spatiotemporally). | [28] |
| Enhancer3<br>*LOC127896430* *NANOG* hESC enhancer | *Alus* act as enhancers and can regulate nearby promoters; *NANOG* is a regulator of human embryonic stem cell (hESC) identity | [29, 30] |

## Results

### Evolution of exonic silencer: Cultivator fixes bases in silencer

The overlapping exonic silencer sequence has dual functions: it encodes the C-terminal end aa sequence of the 68-aa protein and forms a part of the silencer *LOC130067579*. Investigated were how initial bases of the exonic silencer were formed and fixed during the early evolutionary stages of development. We searched for the earliest living species that carries the sequence of *SMIM45.* The available data bank annotations in *Ensembl* show the earliest species with the annotation of *SMIM45* is that of the elephant shark (*Callorhinchus milii*) whose age of divergence is ∼435 Mya. There is no information on earlier animal species that may contain the *SMIM45* locus or its flanking genes, e.g., the sponges, members of the phylum Porifera.

A comparison of the exonic silencer sequence present the elephant shark compared with that of the human sequence shows that twenty-five out of thirty-eight bases (66%) are identical (Fig 2a). These bases were likely fixed in an ancestor to the elephant shark. Thus, much of the exonic silencer sequence is of ancient origins, over 435 Mya. Twenty-four bases are fixed because they are in either the first or second position of a codon (Fig. 2, top).

**Fig.2.**
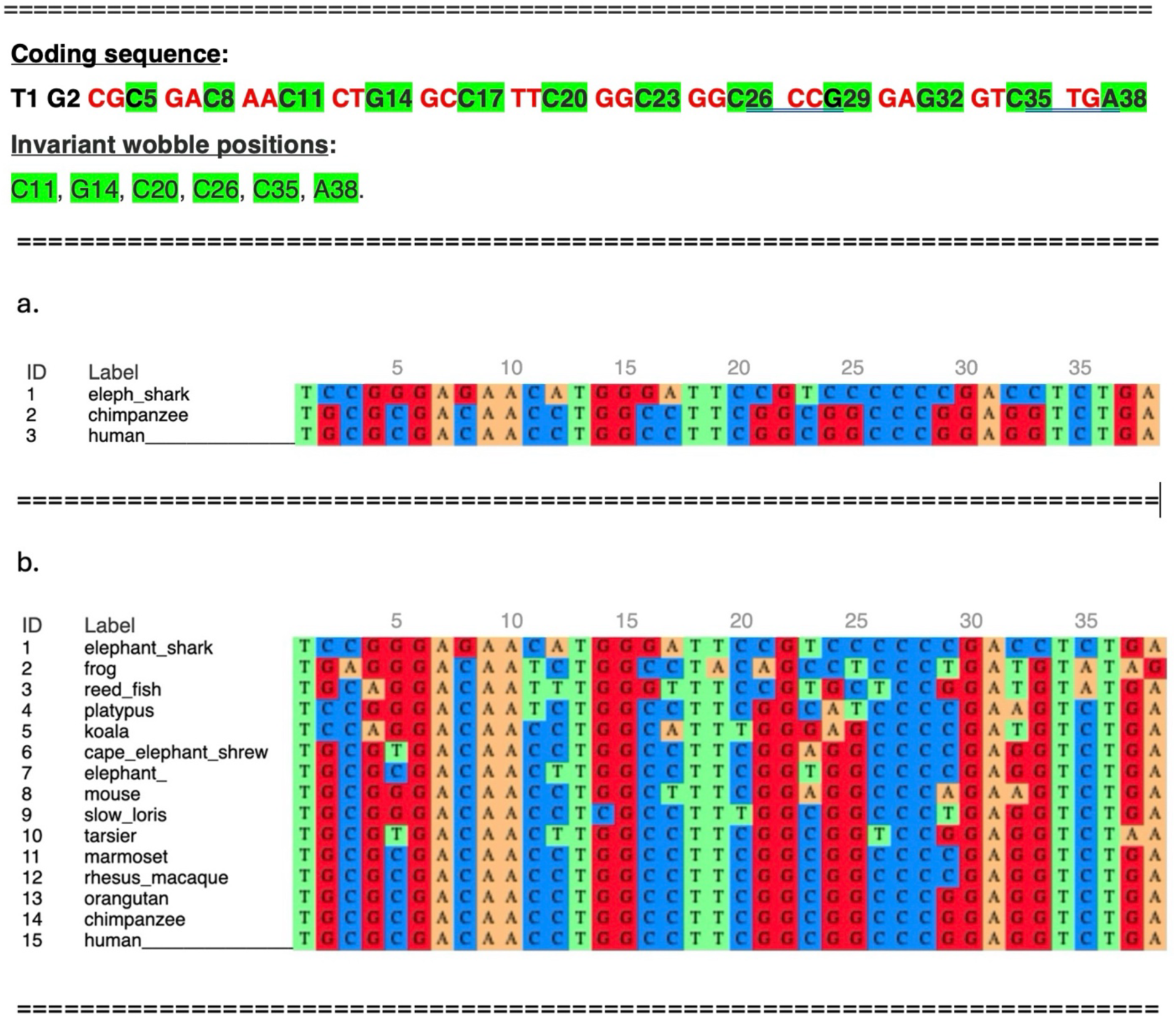
Alignment of the human exonic silencer sequence with sequences from other species. Top. The C-terminal 68-aa coding sequence; highlighted are the first and second codon positions in red, the wobble positions in green. Invariant wobble positions are also shown with green highlight. Fig.2a, alignment of the human and chimp sequences with that of the elephant shark. Fig. 2b. The first codon begins with the third base, CGC. The alignment of base sequences homologous to exonic silencer from ancestral species with that of the sequence of human exonic silencer. The order of alignment of species 1-15 is by age of evolutionary appearance. The alignment is by MAFFT (https://mafft.cbrc.jp/alignment/server/index.html).

A change in bases of codons can result in a change in the aa sequence of the 68-aa C-terminal end. For example, a change in the invariant G6 to C6 translates to the aa sequence RHNLAFGGPEV; the human aa sequence is RDNLAFGGPEV. Thus, twenty-four bases, which also form the sequence of the exonic silencer, are fixed by the cultivator-68 aa sequence and are fixed by natural selection. This is consistent with and supports the cultivator model of Lee et al. [20].

Located within the 38-base silencer/68-aa sequence are twelve wobble bases (Fig. 2, top, highlighted in green). Six of the twelve wobble bases, plus a seventh, T1, which is outside the ORF, are invariant between elephant shark and humans (Fig. 2a and top). Because these DNA bases do not dictate amino acid fidelity, they represent a specialized conservation within the exonic silencer, likely fixed in an ancestor to the elephant shark. This raises the important question of the mechanism of origination and fixation of these seven invariant bases. However, currently this cannot be answered. Sequence analysis of *SMIM45* or its neighbor genes (*CENPM* and *SEPTIN3)* from species ancestral to the elephant shark, such as lamprey, sea star, or sponges, is currently not possible as these genes have not yet been annotated within the available genome assemblies.

Fig. 2b shows the multiple sequence alignment of the 38-base exonic silencer in humans with that of the homologous sequences from fourteen species; it demonstrates that the exonic silencer sequence was evolutionarily completed in the marmoset (a New World primate of the Platyrrhini parvorder), approximately 40 Mya. To determine when the amino-terminal end of the 68-aa ORF aa sequence was completed, 38 nt sequences from different species were translated. Fig. 3 shows that the C-terminal end of the 68 aa ORF is completed in the Afrotheria. Therefore, the completion of the C-terminal end of the 68-aa ORF sequence was in the Afrotheria clade ∼100 Mya, and the exonic silencer sequence was completed in the New World primates ∼40 Mya.

**Fig.3.**
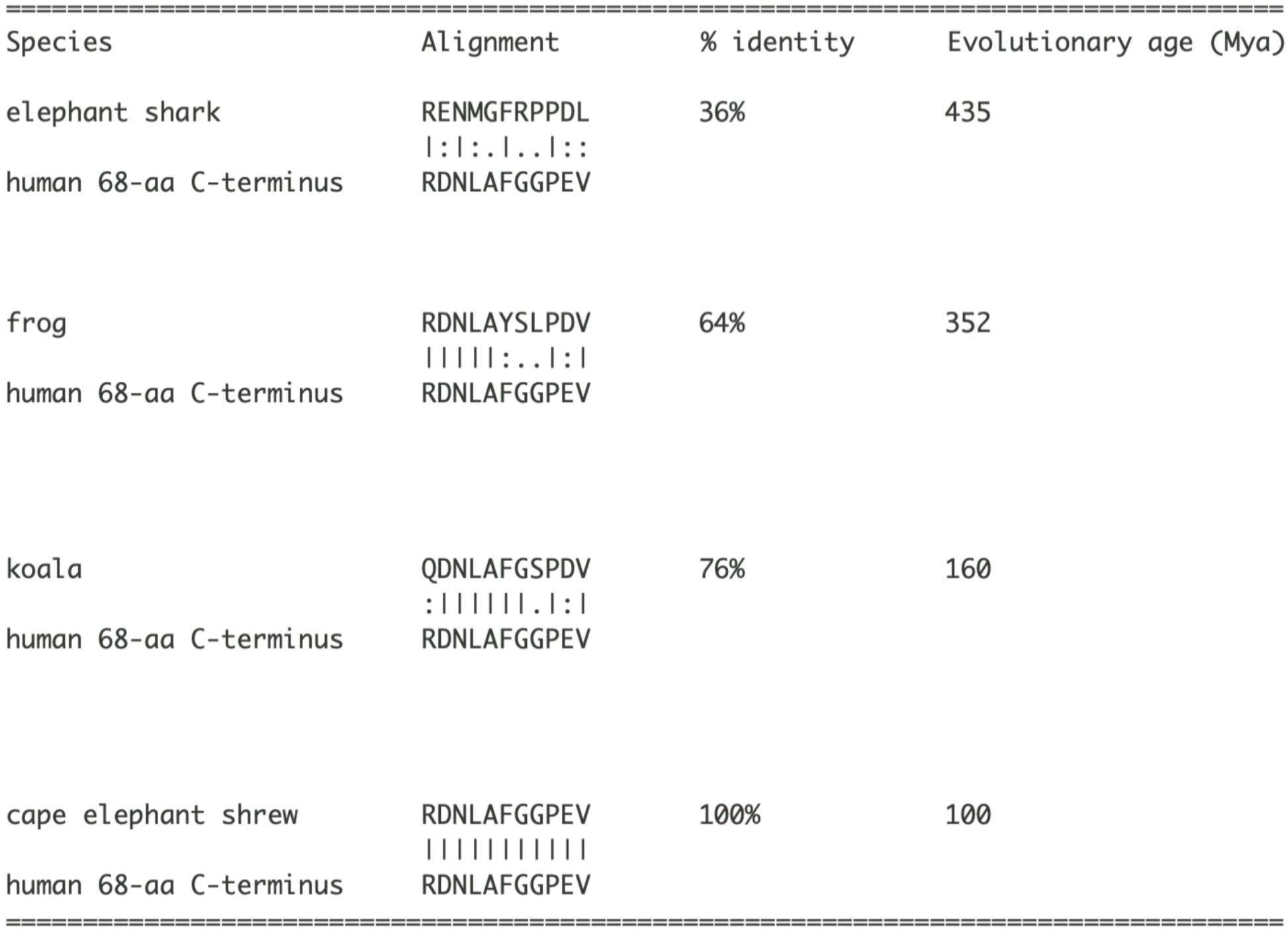
Translated aa sequence from the 38 nt sequence exonic silencer from different species and the percent identities. The <u>SIB Swiss Institute of Bioinformatics</u>, ExPASy Translational tool was used for the translation of nucleotide sequences. The aa sequences represent 5’ 3’ Frame 3 translations; evolutionary ages are for reference. Alignment was with EMBOSS Needle.

The completion of the 68-aa C-terminal sequence within the Afrotheria clade is chronologically linked to the emergence of silencer b, the regulatory segment located in the promoter region. Silencer b was previously shown to have originated in the Afrotheria clade [7]. Given this chronological linkage, we suggest that the fixed/invariant exonic bases perhaps were key in the initial formation of silencer b, though via an unidentified cis-acting mechanism.

### Evolution of exonic silencer: further development via biased mutations at nearest-neighbor invariant bases

We address the evolutionary expansion of the exonic silencer sequence. Analyzed are mutations that occur on the 5’ and 3’ sides of invariant bases. Thirteen bases are considered (Table 2). Excluded are invariant positions in codons where neighbor codon bases are fixed by natural selection. During evolution, nearest-neighbor mutations occurred almost entirely to produce G and C bases (Table 2). The fact that 12 out of 13 neighbors are G and C suggests that invariant bases and the sequence context steer mutational pressure toward GC accumulation within the exonic silencer. In other studies, it has been shown that biased nearest-neighbor mutations that occur during evolution depend on the identity of neighboring bases [32]. Although the small sample size precludes formal statistical analysis, the data show that mutations in nearest neighbors of invariant bases increase the GC and CpG content of the exonic silencer locus. The exonic silencer has a 70% GC content and contains five CpG dinucleotides, with three CpG sites contributed by nearest-neighbor bias. Silencers have been shown to repress gene expression in CpG -rich regions [18]. This may pertain to repression of the expression of 107-aa cistron in somatic tissues. The mutations that occur at neighboring bases of invariant bases result in the total completion of the exonic silencer sequence that is present in humans.

**Table 2.**
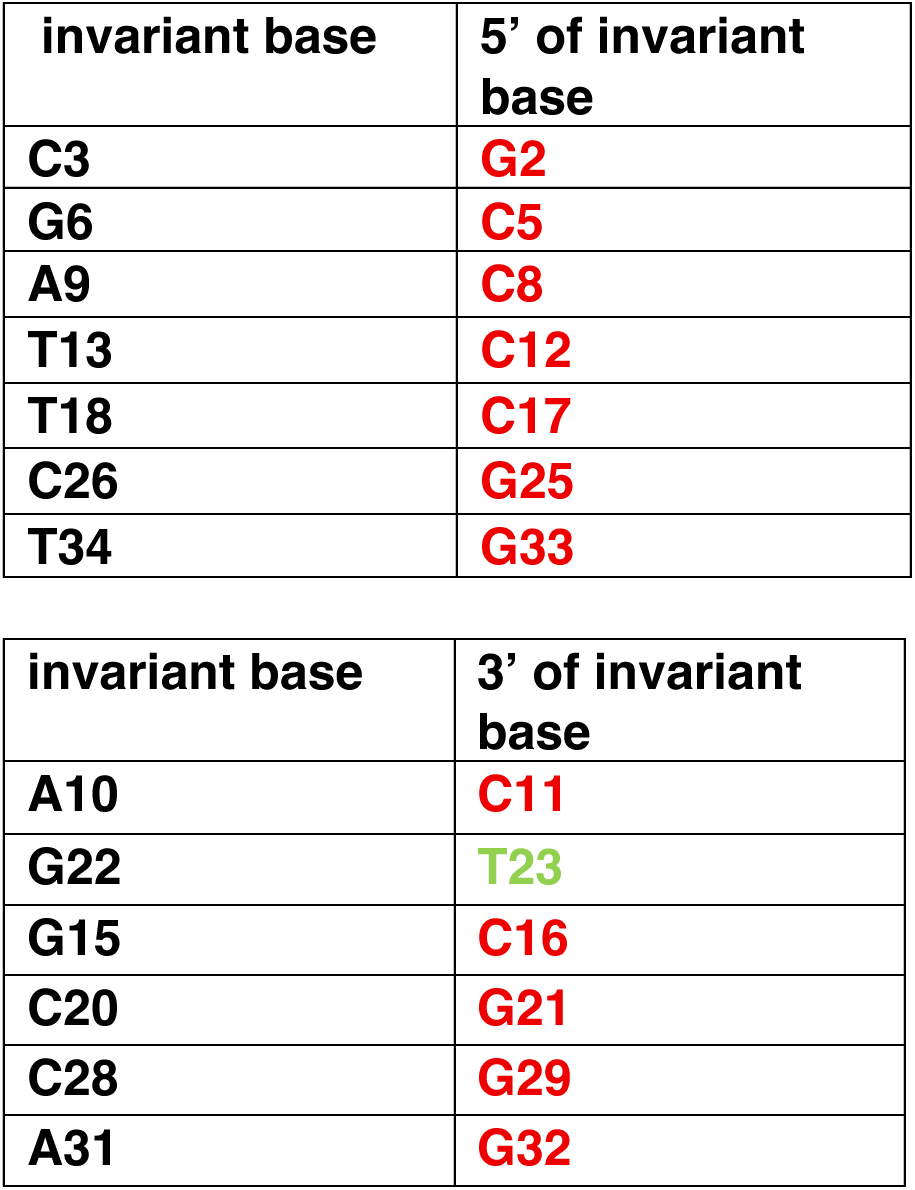
Nearest-Neighbor Bases of Invariant Bases.

| invariant base | 5' of invariant base |
| --- | --- |
| C3 | G2 |
| G6 | C5 |
| A9 | C8 |
| T13 | C12 |
| T18 | C17 |
| C26 | G25 |
| T34 | G33 |

| invariant base | 3' of invariant base |
| --- | --- |
| A10 | C11 |
| G22 | T23 |
| G15 | C16 |
| C20 | G21 |
| C28 | G29 |
| A31 | G32 |

Thus, the exonic silencer is formed through a combination of processes: codon-fixed bases (via cultivator model), GC bias at nearest-neighbor invariant bases, and invariant wobble bases that were formed by unknown means and in unknown ancient species. The invariant wobble bases provide an estimate of over 435 Mya for origination of the exonic silencer. The invariant wobble base analysis updates the previous estimate of ∼352 Mya to ∼429 Mya, which was based on synteny and sequence similarity [7].

### Silencer b (segment of silencer LOC130067579 within the promoter): detection of species-specific sequences

Silencer b originated *de novo* within the Afrotheria clade, by a pathway and at a different time than the exonic silencer [7]. Here we show that the silencer b sequence was completed in the chimpanzee ∼6 Mya (Fig S1). Four point mutations distinguish the chimpanzee/human sequence from that of the gorilla. This contrasts with completion of the exonic silencer in the New World primates ∼40 Mya. Although the exonic silencer and silencer b differ significantly in their mechanisms and timeframe of origin, both constitute the sequence of silencer *LOC13006757* and function within this context. As silencer b originated *de novo*, we searched for a proto-silencer b sequence. Fig.4 shows an alignment of silencer b with orthologous sequences from the Afrotheria and the Great Apes. The longest sequence found to be invariant is cctctgcagcc (Fig. 4). This 11-base sequence is 100% conserved across the deeply divergent mammalian lineages (Afrotheria to Primates, approximately 90 million years of separation). Furthermore, there is a synteny with the 68-aa bp sequence. Such high levels of conservation point to strong purifying selection and suggest that this sequence is likely a proto-silencer. As mentioned before, the fixed/invariant bases within the exonic silencer might have served a role by facilitating the formation of silencer b, with the necessary information for this process stored directly within those bases. However, how fixed exonic silencer bases may have initiated the proto-silencer b sequence is unknown, despite their suggested importance that was mentioned previously. For reference purposes, the Afrotheria phylogeny is shown in Fig. 5.

**Fig.4.**
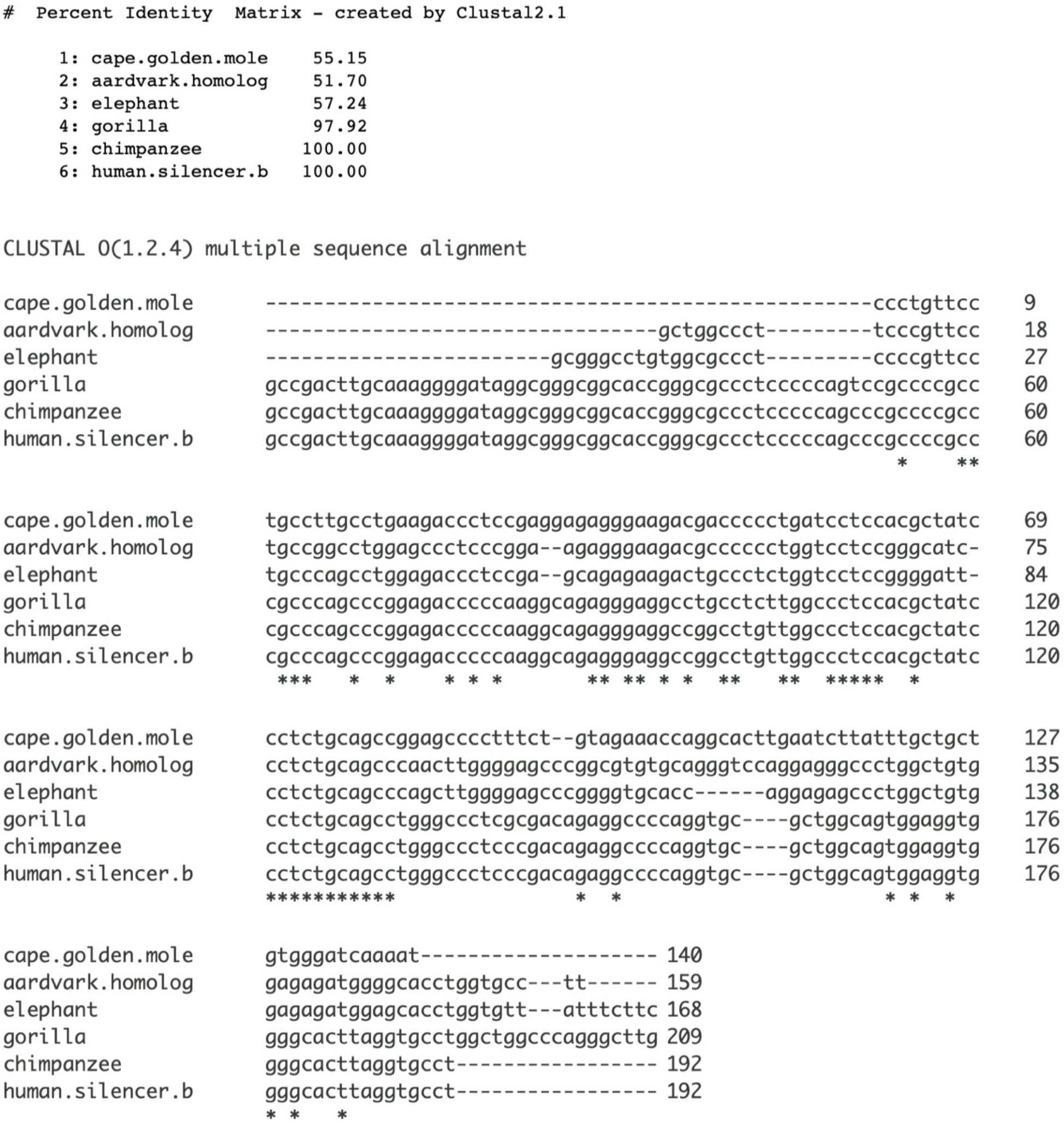
Alignment of silencer b with homologous sequences from representative Afrotheria and Great Apes species. Base positions 121-131 are totally conserved. Clustal Omega Multiple Sequence Alignment (MSA) was used.

**Fig. 5.**
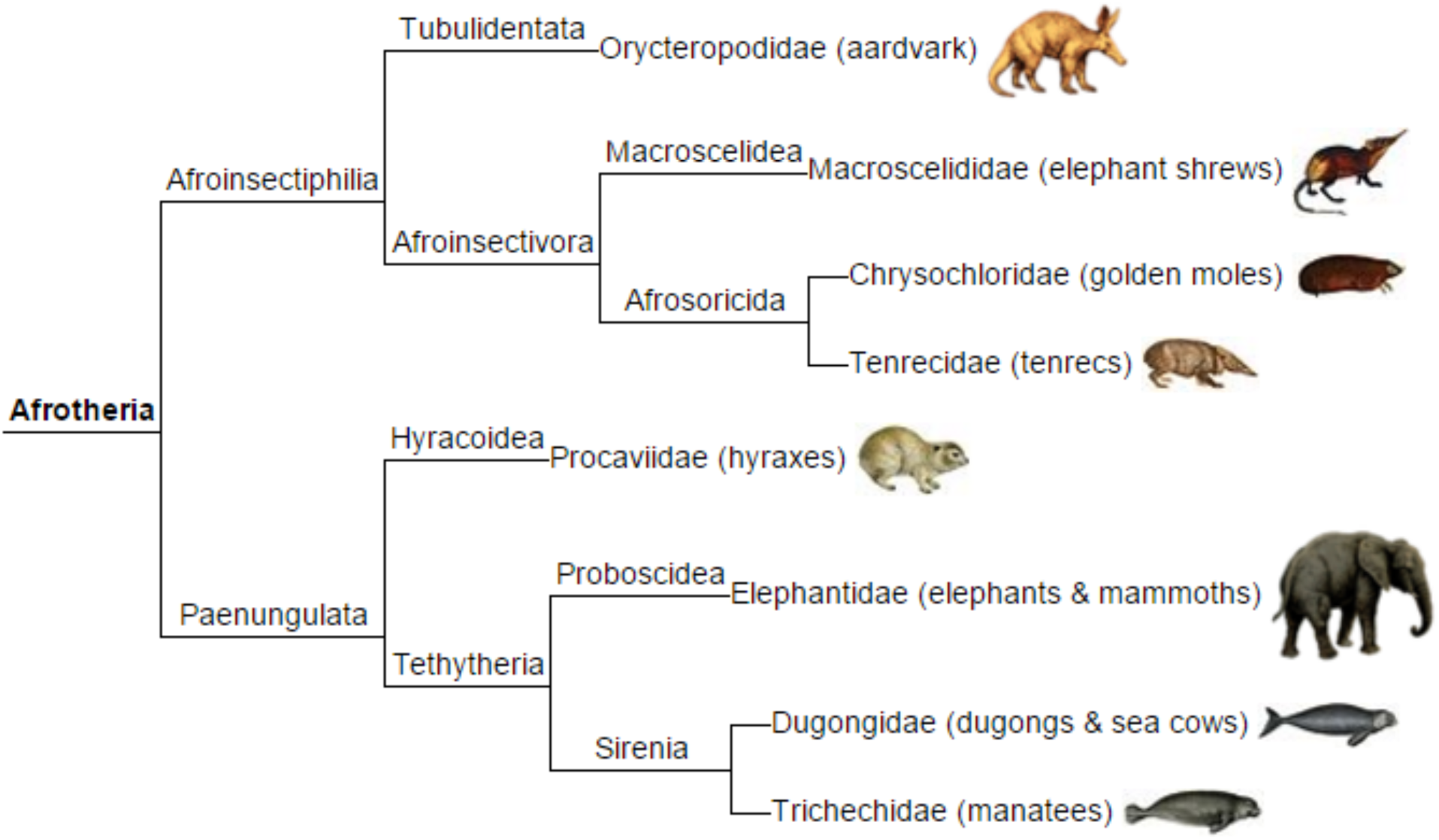
Phylogeny of the Afrotheria clade. Illustration is from WikiCommons and is in the public domain.

Alignment of the silencer sequence with homologs from Afrotheria, Old World monkeys, and Great Apes reveals that a repeat, gcccgcccc was formed in the Great Ape *SMIM45* locus during evolution (Fig. S2). It is part of a longer repeat: gcccgccccgcccgccc. This indicates evolutionary pressure to increase the GC and CpG content of silencer b in the Great Apes. Silencer b contains thirteen copies of CpG. Together with the five CpG dinucleotides present in the exonic silencer, there are eighteen copies of CpG in silencer *LOC130067579,* ATACSTARR-seq lymphoblastoid silent region 13815. This is consistent with the repression of gene expression that is dependent on high CpG content of a locus [17,18].

### Enhancer1: sequence origin and completion

Enhancer 1 is situated eight base pairs from the 5’ start of the *SMIM45* gene and also resides within the exons of four transcript variants. Here, we investigate the evolutionary origin and sequence completion of the enhancer. Alignment with homologous regions in other primate species and in close primate relatives suggests that enhancer 1 originated *de novo*. The lemur sequence displays significant similarity with enhancer 1 (78%), the tree shrew, also a close relative to the primates, shows no significant identity (40%) (Fig. 6). The age of divergence from a common ancestor is ∼63 Mya for the lemur and ∼68 Mya for the tree shrew. The data suggest that enhancer 1 originated in an extinct early, stem-lineage of the lemur more than 60 Mya. It was not possible to identify a putative proto-enhancer 1 as sequence alignments with the tree shrew could not be made with accurate synteny. For reference, a phylogenetic tree of the primates is in Fig. S3.

**Fig.6.**
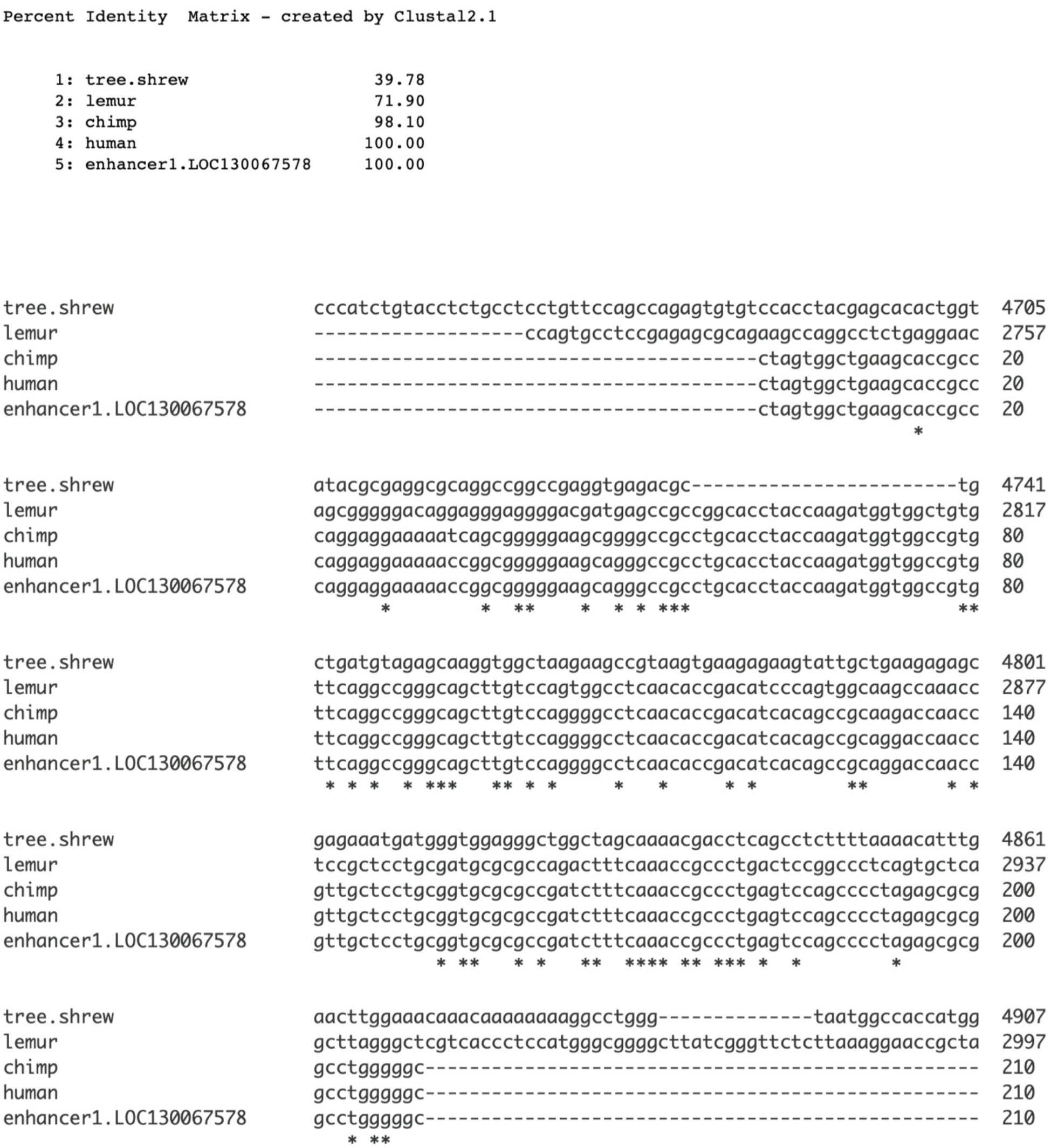
Alignment of enhancer 1 with genomic sequences of the tree shrew (*Tupaia chinensis*, Chinese tree shrew) and the lemur (*Lemur catta*, Ring-tailed lemur). Both species are the closest living relatives to the primates. Clustal Omega Multiple Sequence Alignment (MSA) was used for alignment.

Additionally, the species for which the enhancer 1 sequence was finalized was investigated. Fig. 7 shows there are four-point mutations that distinguish the human from the two chimpanzee sequences. Enhancer 1 was thus completed in the human genome.

**Fig.7.**
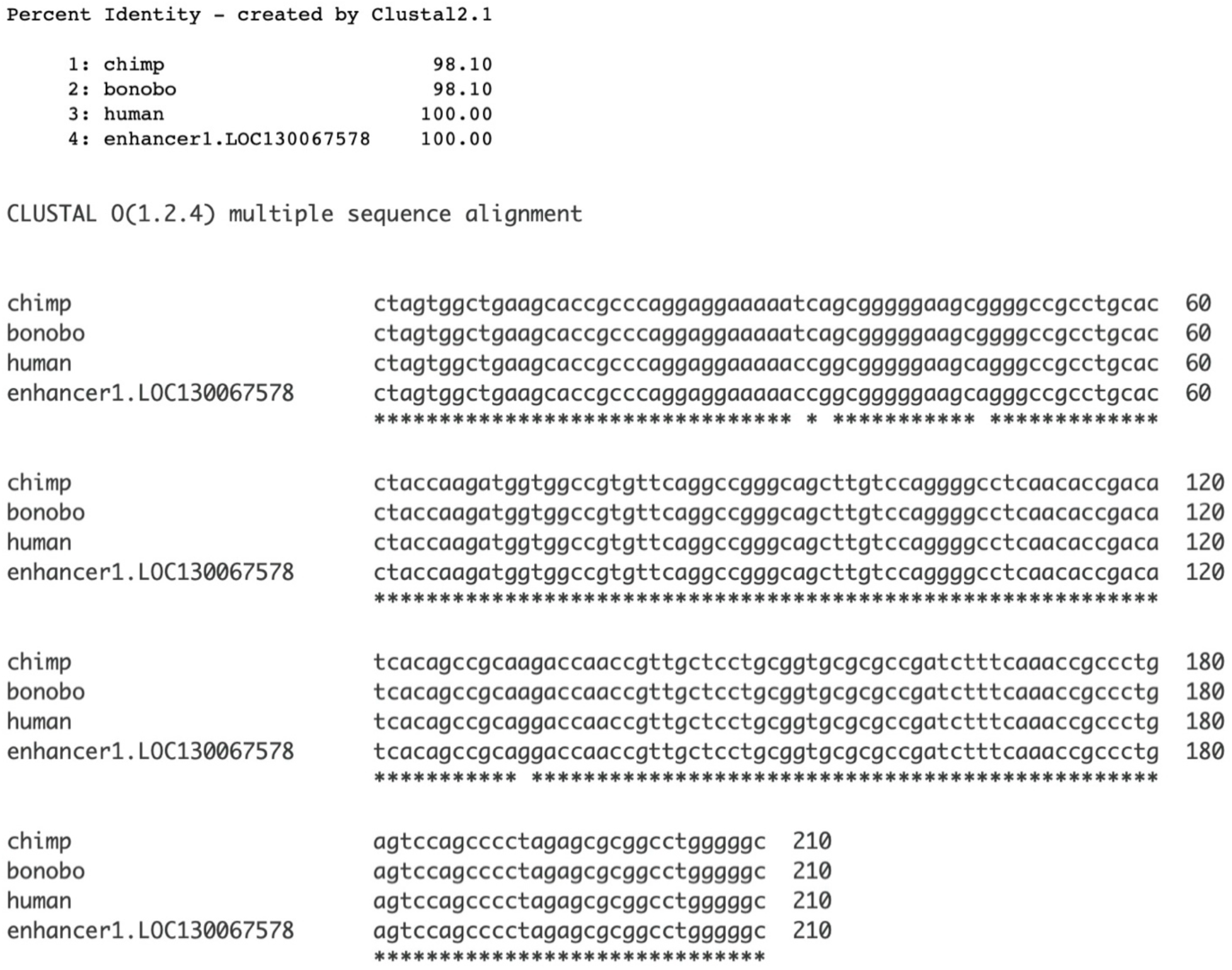
Completion of enhancer 1 sequence in humans. Clustal Omega Multiple Sequence Alignment (MSA) was used was used for alignment.

### Enhancer2

The evolutionary origination and completion of enhancer 2, *LOC127896429* H3K4me1hesc was investigated. Similar to that of enhancer 1, enhancer 2 originated by the *de novo* process. Sequences from five members of the Afrotheria clade were aligned with those of the chimpanzee and the human enhancer 2. The percent identities show that sequences from several Afrotheria species approach the “twilight zone” of significance, suggesting origination from the Afrotheria clade (Fig. 8, top). The data suggest that enhancer 2 originated within the Afroinsectivora branch of the Afrotheria clade (Fig. 5). The koala (*Phascolarctos cinereus*) lineage separated from the ancestor of Afrotheria before the Afrotheria separated from other placental mammals. Its genomic sequence does not significantly align with enhancer 2 (45% identity). This is consistent with the determination that enhancer 2 originated in the Afrotheria clade, ∼100 Mya. One segment of enhancer 2 displays invariance, 17cagtcac23 with exception of the tenrec sequence (Fig. 8, bottom); this sequence may be a candidate for a proto-enhancer 2 as the 8-base sequence is under purifying selection, over ∼90 millions of years of separation.

**Fig. 8.**
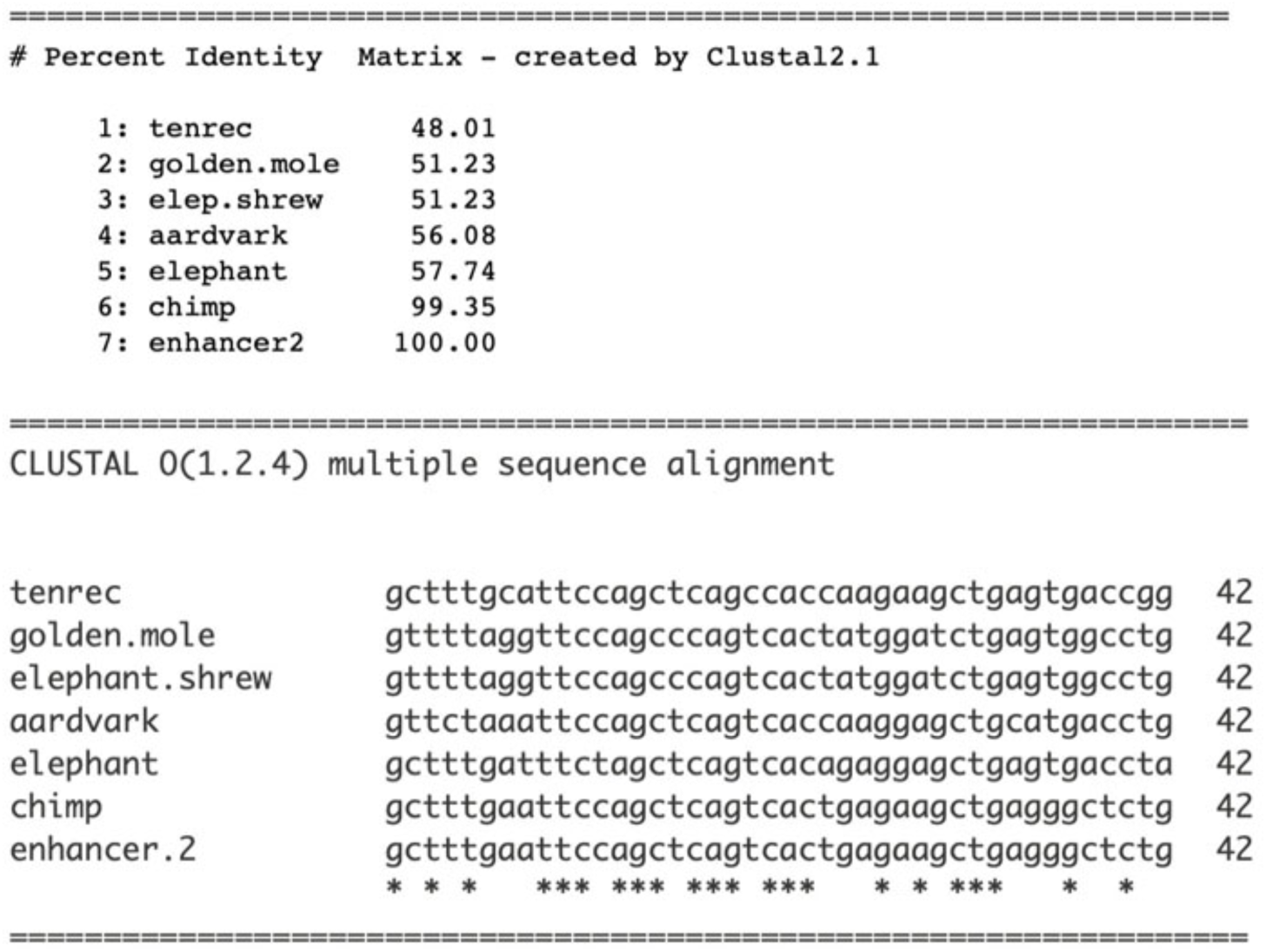
Homologous sequences from the Afrotheria clade were aligned with sequences of the chimpanzee and enhancer 2. Top. The percent identities. Bottom, a segment of enhancer 2 aligned with homologous sequences Clustal Omega Multiple Sequence Alignment (MSA) was used.

An alignment of the chimpanzee and human *SMIM45* sequences with that of the enhancer 2 reveals two bp deletions and three point mutations that distinguish the human sequence from that of the chimpanzee (Fig. S4). Therefore, this identifies enhancer 2 as human specific. Nevertheless, similar to the mutations in enhancer 1, the functional impact of these mutations remains unknown. The completion of the embedded silencer, *LOC127896429* silent region_13814 was also determined. The sequence was completed in the chimpanzee (Fig.S5) and therefore it is not human specific. Interestingly, this contrasts with the completion of enhancer 2.

### Enhancer 3, NANOG hESC: birth by Alu insertions

Branco and Chuong [33] provide examples of how transposable elements can drive regulatory innovation. Functioning in gene regulation, many enhancers contain *Alu* transposable elements that modulate target gene expression through promoter interactions [34–36]. Nearly half of *Alu* elements present in the human genome are in introns where they regulate gene transcription in specific tissues [37]. In accordance with this, enhancer 3 (*LOC127896430* NANOG hESC) contains two *Alu* elements (Fig. 9). In *SMIM45* RNA transcripts, enhancer 3 resides in the intron of *SMIM45* RNA transcript variant 1, NM_001395940. Given its intron location, perhaps enhancer 3 may participate in the regulation of the 107-aa cistron in embryonic brain tissues.

**Fig.9.**
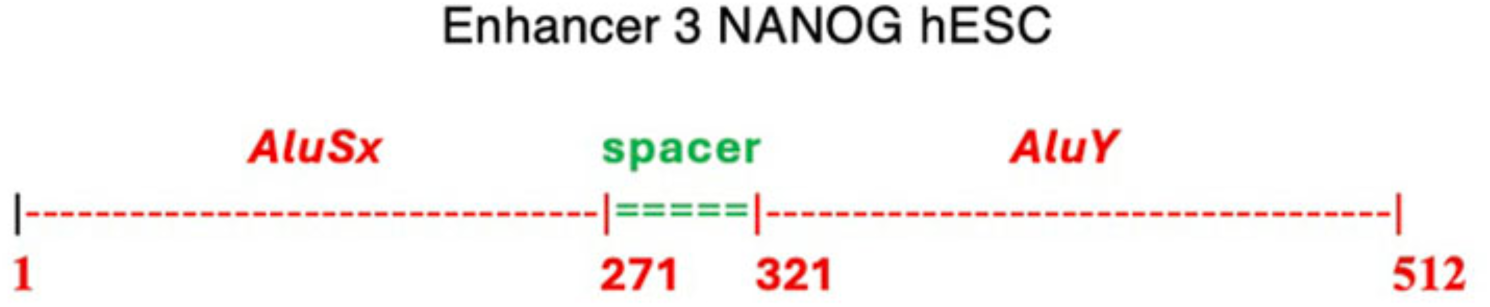
Schematic of components of enhancer 3. *Alu* TEs and the spacer sizes were determined by RepeatMasker [38].

Enhancer 3 comprises tandem transposable elements (TE) *AluSx* and *AluY* that are separated by a 50 bp spacer (Fig. 9). To investigate evolutionary origins, *SMIM45* orthologs from various primate species were aligned with the human enhancer 3. Fig. 10 demonstrates that the enhancer originated in the Old World monkey lineage, as evidenced by its presence in the rhesus monkey (*Macaca mulatta*) and olive baboon (*Papio 20nubis*) *SMIM45* loci. The enhancer sequence was not detected in Ma’s night monkey (*Aotus nancymaae*), a New World primate. Both the rhesus monkey and olive baboon diverged from the human lineage approximately 25-30 million years ago. Fig. 10 shows that the rhesus and baboon sequences possess the complete *AluSx* sequence along with part of the spacer sequence. The enhancer appears to have originated by the insertion of the *AluSx* TE into the Old World monkey (*Catarrhini*) *SMIM45* locus. A second *Alu*, *AluY* was inserted in the Great Apes (*Hominoidae*) *SMIM45* as it is found only in the orangutan and the other Great Apes. For reference, Fig.S3 provides a diagram of primate phylogeny.

**Fig. 10.**
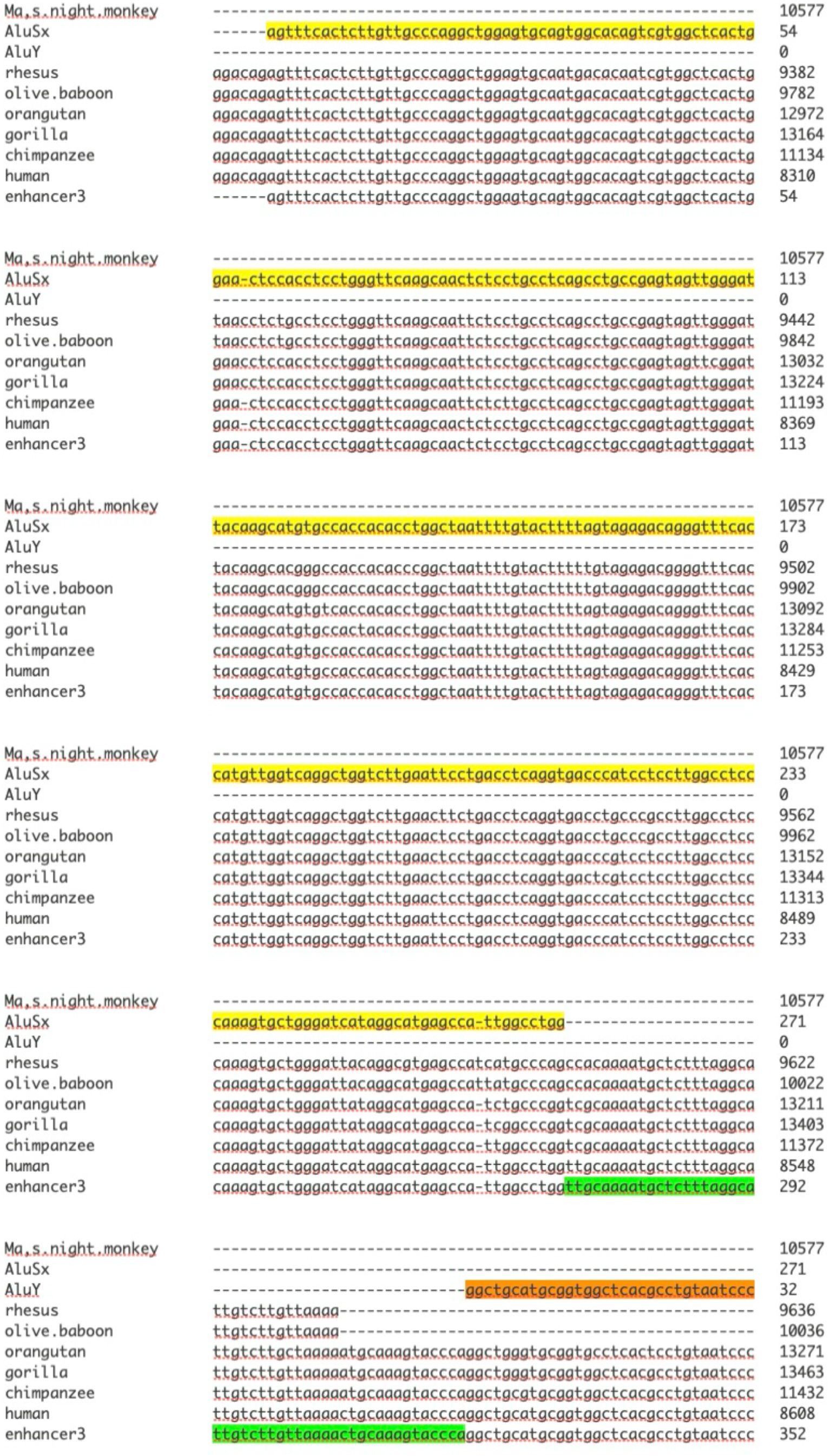
Alignment of homologous primate species sequences with that of enhancer 3. Highlighted regions indicate *AluSx* (yellow), *AluY* (brown), and the intervening spacer (green). Note the alignment with the *AluY* does not show the total alignment.

To determine in which species the enhancer 3 sequence was completed, we aligned the chimpanzee sequence with the human enhancer 3 sequence (Fig. S6). Twelve point mutations differentiate the human and chimp/bonobo sequences from enhancer 3. Thus, the enhancer 3 sequence is human specific. Enhancer 3 contains the sequence TAATTTTGT, which represents the transcription factor NANOG core binding motif consisting of TAAT followed by several Ts and GT [39].

## Discussion

This study investigates the evolutionary origins of regulatory sequences within the *SMIM45* locus, providing insights into the mechanisms underlying the formation of silencers and enhancers. Notably, we show that over a massive time span of >400 My, *SMIM45* evolved a suite of regulatory elements believed to control the human specific 107-aa protein’s activity during human fetal brain development. The complexity of these combined regulatory processes that are vital for human embryonic brain development may explain this long evolutionary time frame. The exonic silencer sequence, which overlaps the 68-aa C-terminal end, is the first of the regulatory sequences to originate evolutionarily, featuring a small number of anciently formed invariant wobble bases. This invariance reflects the silencer’s functional constraints. While the mechanism of fixation of these bases is unknown, the majority of exonic silencer bases are fixed by the cultivator model described by Lee et al [20]. The data show that the cultivator-fixed bases and the wobble invariant bases carry information that guides exonic silencer maturation via nearest neighbor bias, resulting in the total completion of the 38-base-pair exonic silencer. *De novo* origination of both silencer b (segment of the ATAC-STARR-seq lymphoblastoid region 13815 in the promoter region) and the 107-aa protein cistron occurred in the Afrotheria clade [7]. Concurrently, the 68-aa C-terminal base sequence was also completed in Afrotheria, hinting at a potential role of these bases (that also make up the exonic silencer-specific fixed bases) in the emergence of Silencer b. Pang and Snyder reported that a large number of silencers are found in exons of genes [23]; however, our literature search yielded no existing presence of exonic silencers that partially overlap ORFs, with the exception of our previous study [7]. The ATAC-STARR-seq lymphoblastoid silent region 13815 within the *SMIM45* gene appears to be the first of its kind to be identified and characterized, and is thus a novel regulatory element.

Pu *et al* [40] provided a detailed study on the evolution of enhancers containing embedded silencers. By analyzing expression patterns across *Drosophila melanogaster* and related species, they demonstrated that the evolution of repressor sequences can precede that of the enhancer sequences. Although the embedded silencer in the enhancer 2 sequence reported here was completed before the enhancer, the significance of this remains uncertain due to the short evolutionary time span.

We have demonstrated that bias in nearest neighbors of invariant bases in the exonic silencer raises the GC content of the locus to 73% during evolution. Similarly, there is evolutionary pressure to increase GC content within silencer b, which is in the promoter of the 107-aa cistron. High GC content facilitates chromatin accessibility, enabling the transcriptional apparatus to access DNA [41, 42] and potentially mediating suppression. We also demonstrate evolutionary pressure to add the CpG motif to the silencer *LOC130067579* sequence. There is a known role of high CpG content in gene repression [18]. Yang *et al* [43] confirmed that CpG content impacts gene expression evolution and silencers are recognized for repressing somatic gene expression [44]. The presence of a high GC and CpG content in *LOC130067579* is consistent with the concept that this silencer suppresses the 107-aa cistron in somatic tissues.

The completion of all three enhancer sequences in the human *SMIM45* locus points to the regulation of the human 107-aa cistron, especially given that the orthologous chimpanzee sequence encodes a truncated protein [7]. Concerning *SMIM45* regulatory element functions, a comparison with known related silencers and enhancers is instructive. Although it is not a bicistronic gene, *FGF18* (fibroblast growth factor 18) is similar to *SMIM45* in having super-silencers and super-enhancers [45]. Two silencers in the *FGF18* gene can act together through compensatory chromatin interactions. Perhaps compensatory chromatin interactions also relate to the function of the two silencers in *SMIM45, silent* region_13814 and silent_region 13815.

Both Enhancer 1 and enhancer 2 originated *de novo* from ancestral non-coding regions and appear to be part of a large class de novo of human brain specific enhancers [46]. These brain-specific enhancer sequences are completed by one or a small number of point mutations distinguishing human from chimpanzee sequences. Consistent with this, enhancer 1 and enhancer 2 show a small number of mutations that complete the human sequence. *De novo* enhancers can induce genes that are critical to cognitive function and are expressed in progenitor cells of the developing neocortex [46]. By analogy, enhancers 1 and 2 may regulate the expression of the *SMIM45* 107-aa protein shown to be produced in the embryonic cerebral cortex [8].

Enhancer 3, LOC127896430 NANOG hESC enhancer consists of two *Alu* elements and contains the transcription factor NANOG core binding motif. NANOG is known to bind enhancers in embryonic stem cells to regulate gene expression during developmental processes [47, 48]. This may relate to the regulation of the 107-aa cistron by enhancer 3 as the 107-aa protein is expressed in embryonic brain tissues [8].

We have considered the regulatory elements individually, however, *SMIM45* contains a complex of super-enhancer and super-silencer. These act similarly to the individual elements except function in an intense fashion; super-enhancers to strongly boost gene expression in a cooperative manner; super-silencers to act collectively and exert strong, coordinated suppression of gene expression.

The aforementioned analogies provide important insights as to how they may activate or suppress the 107-aa cistron. These enhancers and silencers remain promising targets for future studies.

## Conclusions

*SMIM45* appears to represent a unique bicistronic gene characterized by an intricate suite of regulatory sequences. Evolving over several hundred million years, these transcriptional enhancers and silencers emerged through complex, distinct processes. Silencer *LOC130067579,* ATACSTARR-seq lymphoblastoid silent region consists of two segments, one overlapping an ORF (exonic silencer) and the other overlapping the 107-aa promoter (silencer b). These two segments developed independently across different evolutionary timeframes; notably, the exonic silencer formed through diverse processes and was the first regulatory sequence to originate during evolution. The results point to a complex regulation of the *de novo* 107-aa cistron during human embryonic brain development.

## Methods

### Source of properties of human SMIM45

The NCBI Gene database (https://www.ncbi.nlm.nih.gov/gene) is the source for *SMIM45* properties, including enhancers, silencers and their experimental evidence, accessible chromatin regions, functions, sequences, and RNA transcript exon/intron sequences.

### SMIM45 in other species

The *SMIM45* orthologous sequences from various species were also obtained from the NCBI Gene database (https://www.ncbi.nlm.nih.gov/gene) except for that of the elephant shark. The elephant shark *SMIM45* sequence is present on the website: *Ensembl:* http://may2025.archive.ensembl.org/Alignment Methods. For species where *SMIM45* has not been annotated, the total sequence between flanking genes *CENPM* and *SEPTIN 3* was used for alignments.

### Nucleotide and amino acid sequence alignments

Clustal Omega, Multiple Sequence Alignment and Pairwise Sequence Alignment EMBOSS Needle Tools < EMBL-EBI) [49] were used.

Several sequences were aligned using MAFFT [50]. (https://mafft.cbrc.jp/alignment/server/index.html). This sequence program allowed for easy visualization of conserved bases.

### CpG pairs in silencer b (in the promoter)

There are 13 CpG pairs in silencer b. The probability of finding 13 Cs followed by 13 Gs in a sequence of 192 bases containing 73% G+C is approximately 6.956×10^-10^ (determined by Google statistical analysis). This supports the statement of a high CpG content present in silencer b.

### Sequences of silencer LOC130067579 ATAC-STARR-seq lymphoblastoid silent region 13815 and its segments

Silencer *LOC130067579*

tgcgcgacaacctggccttcggcggcccggaggtctgagccgacttgcaaaggggatagg cgggcggcaccgggcgccctcccccagcccgccccgcccgcccagcccggagacccccaa ggcagagggaggccggcctgttggccctccacgctatccctctgcagcctgggccctccc gacagaggccccaggtgcgctggcagtggaggtggggcacttaggtgcct 230 bps

exonic silencer (overlap with 68-aa)

tgcgcgacaacctggccttcggcggcccggaggtctga 38 bps

silencer b (in the promoter)

gccgacttgcaaaggggataggcgggcggcaccgggcgccctcccccagcccgccccgcc cgcccagcccggagacccccaaggcagagggaggccggcctgttggccctccacgctatc cctctgcagcctgggccctcccgacagaggccccaggtgcgctggcagtggaggtggggc acttaggtgcct 192 bps

#### Enhancer sequences

Enhancer 1, *LOC130067578* ATAC-STARR-seq lymphoblastoid active region 19151 [*Homo sapiens* (human)] Gene ID: 130067578

ctagtggctgaagcaccgcccaggaggaaaaaccggcgggggaagcagggccgcctgcacctaccaagatggtggccgtgttcaggccgggcagcttgtccaggggcctcaacaccgacatcacagccgcaggaccaaccgttgctcctgcggtgcgcgccgatctttcaaaccgccctgagtccagcccctagagcgcggcctgggggc 210 bps

Enhancer 2 *LOC127896429* H3K4me1 hESC enhancer GRCh37_chr22:42346983-42347610 [*Homo sapiens* (human)] Gene ID: 127896429

gagactccgtctcaaaaaaaacaaaccctctgtgaactcacagtcaccccccagtcccacatatgctggaaaggacctg tcatacctgaagagcccctagatggcgcagaggtgtctgtggtgggggacctaggtcctgaagccacctcacccagagg ctttccccctgcccatccccaggtttctgggaacggattccctagggaggtggttcctggaagccttttcccagccacgcccc gtgggccctagggggctgctctctccctcctgagaatagccctcaacacgtggcagataccttgtctatggcataggggga gggggaggatccatgcttgggaaggtggaccccacccccaacgtcagctcttggctttgaattccagctcagtcactgag aagctgagggctctgggagaaggagaaggccagcagcatcacctctctgcctcatcccaaaatggggtctcaacacca atccagctgggaggactgcaggaagtgatgttggggccagctggaagatgggagtgctcaatgcctgtgctggctgtaca ccagccaggggtgctgtggggtagatgaggcagaatggggagggggagccatttgcaagggtcctgaa 628 bps

Embedded silencer in LOC127896429 (silent region_13814)

ctagatggcgcagaggtgtctgtggtgggggacctaggtcctgaagccacctcacccaga 60 bps

Enhancer 3 LOC127896430 NANOG hESC enhancer GRCh37_chr22:42351209-42351720 [Homo sapiens (human)] Gene ID:

127896430agtttcactcttgttgcccaggctggagtgcagtggcacagtcgtggctcactggaactccacctcctgggtt caagcaactctcctgcctcagcctgccgagtagttgggattacaagcatgtgccaccacacctggctaattttgtacttttagt agagacagggtttcaccatgttggtcaggctggtcttgaattcctgacctcaggtgacccatcctccttggcctcccaaagtg ctgggatcataggcatgagccattggcctggttgcaaaatgctctttaggcattgtcttgttaaaactgcaaagtacccaggc tgcatgcggtggctcacgcctgtaatcccagcactttgggaggccgaggtgggcggatcacgaggtcaggagatcaaga ccatcctggctaacacggtgaaaccccgtctctactaaaaatacaaaaaattagctgggtgcagtggcggtcacctgcag tcccagctactcaggaggctgaggcaggag 512 bps

### Determinations of Alus in enhancer 3

RepeatMasker [38] enabled the determination off the *Alu* transposable elements and the location in *SMIM45*.

### Translation of nucleotide sequence

The ExPASy Translational tool website https://web.expasy.org/translate/ [51] was employed to translate the 68-aa C-terminal end nucleotide sequence.

## Supporting Information

Supplementary File S1

