## Supplementary File S1 for "Diverse processes drive the origination and maturation of super-enhancers/super-silencers during a vast evolutionary timescale of a bicistronic gene"

### Diverse processes drive the origination and maturation of super-enhancers and super-silencers during a vast evolutionary timescale of the bicistronic gene

Nicholas Delihas
Department of Microbiology and Immunology
Renaissance School of Medicine; Stony Brook University; Stony Brook, NY 11794 USA
Orchid https://orcid.org/0000-0002-1704-2587

Fig. S1. Alignment of human silencer b with homologous sequences from the gorilla and chimpanzee. While the chimpanzee *SMIM45* gene contains a silencer b sequence identical to that of humans, both species show three point mutations compared with the homologous gorilla sequence. Clustal Omega, Multiple Sequence Alignment was used for alignment.

```
CLUSTAL O(1.2.4) multiple sequence alignment


gorilla              gccgacttgcaaaggggataggcgggcggcaccgggcgccctcccccagtccgccccgcc      60
chimpanzee           gccgacttgcaaaggggataggcgggcggcaccgggcgccctcccccagcccgccccgcc      60
human.silencer.b     gccgacttgcaaaggggataggcgggcggcaccgggcgccctcccccagcccgccccgcc      60
                     ************************************************* **********

gorilla              cgcccagcccggagacccccaaggcagagggaggcctgcctcttggccctccacgctatc      120
chimpanzee           cgcccagcccggagacccccaaggcagagggaggccggcctgttggccctccacgctatc      120
human.silencer.b     cgcccagcccggagacccccaaggcagagggaggccggcctgttggccctccacgctatc      120
                     ************************************ **** ******************

gorilla              cctctgcagcctgggccctcgcgacagaggccccaggtgcgctggcagtggaggtggggc      180
chimpanzee           cctctgcagcctgggccctcccgacagaggccccaggtgcgctggcagtggaggtggggc      180
human.silencer.b     cctctgcagcctgggccctcccgacagaggccccaggtgcgctggcagtggaggtggggc      180
                     ******************** ***************************************

gorilla              acttaggtgcctggctggcccagggcttg     209
chimpanzee           acttaggtgcct-----------------     192
human.silencer.b     acttaggtgcct-----------------     192
                     ************
```

Fig.S2. Alignment diagram shows the GC sequence addition in the Great Apes, except for that of the orangutan. GC addition in alignment silencer b with species of the Afrothere and primates. MAFFT (https://mafft.cbrc.jp/alignment/server/index.html) was used for alignment.

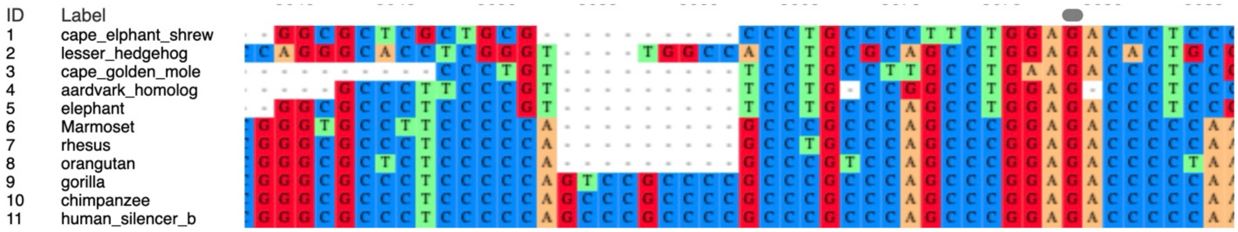


Fig. S3. Diagram of the primate phylogenetic tree. Drawing is from Wikimedia Commons and is in the public domain.

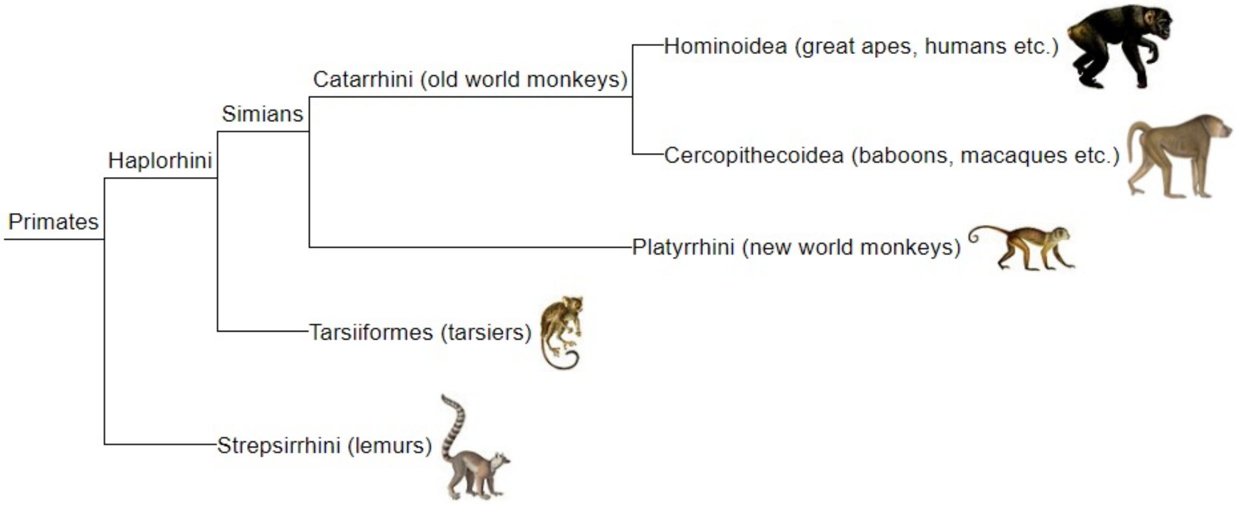

Fig. S4. The alignment of the chimpanzee sequence homologous to enhancer 2 with the *SMIM45* enhancer 2. Alignment shows two bp deletions and three point mutations that distinguish the human sequence from that of the chimpanzee. Clustal Omega, Multiple Sequence Alignment was used for alignment.

```
chimpanzee      ccagcctggcgaacacagcgagactccgtctcaaaaaaaacaaaccctctgtgaactcac 6900
enhancer2       -------------------gagactccgtctcaaaaaaaacaaaccctctgtgaactcac 41
                                   *****************************************

chimpanzee      agtcacccccagtcccacatatgctggaaaggacccgtcatacctgaagagcccctaga 6960
enhancer2       agtcacccccagtcccacatatgctggaaaggacctgtcatacctgaagagcccctaga 101
                *********************************** ************************

chimpanzee      tggcgcagaggtgtctgtggtgggggacctaggtcctgaagccacctcacccagaggctt 7020
enhancer2       tggcgcagaggtgtctgtggtgggggacctaggtcctgaagccacctcacccagaggctt 161
                ************************************************************

chimpanzee      tcccccctgcccatccccaggtttctgggaacggattccctagggaggtggttcctggaag 7080
enhancer2       tcccccctgcccatccccaggtttctgggaacggattccctagggaggtggttcctggaag 221
                ************************************************************

chimpanzee      cctcttcccagccacgccctgtgggccctagggggctgctctctccctcctgagacaata 7140
enhancer2       ccttttcccagccacgccccgtgggccctagggggctgctctctccctcctga--gaata 279
                *** *************** *********************************   ****

chimpanzee      gccctcaacacgtggcagataccttgtctatggcatagggggagggggaggatccatgct 7200
enhancer2       gccctcaacacgtggcagataccttgtctatggcatagggggagggggaggatccatgct 339
                ************************************************************

chimpanzee      tgggaaggtggaccccacccccaacgtcagctcttggctttgaattccagctcagtcact 7260
enhancer2       tgggaaggtggaccccacccccaacgtcagctcttggctttgaattccagctcagtcact 399
                ************************************************************

chimpanzee      gagaagctgagggctctgggagaaggagaaggccagcagcatcacctctctgcctcatcc 7320
enhancer2       gagaagctgagggctctgggagaaggagaaggccagcagcatcacctctctgcctcatcc 459
                ************************************************************

chimpanzee      caaaatggggtctcaacaccaatccagctgggaggactgcaggaagtgatgttggggcca 7380
enhancer2       caaaatggggtctcaacaccaatccagctgggaggactgcaggaagtgatgttggggcca 519
                ************************************************************

chimpanzee      gctggaagatgggagtgctcaatgcctgtgctggctgtacaccagccaggggtgctgtgg 7440
enhancer2       gctggaagatgggagtgctcaatgcctgtgctggctgtacaccagccaggggtgctgtgg 579
                ************************************************************

chimpanzee      ggtagatgaggcagaatggggagggggagccatttgcaagggtcctgaaagccaggctag 7500
enhancer2       ggtagatgaggcagaatggggagggggagccatttgcaagggtcctgaa----------- 628
                *************************************************
```

Fig. S5. The evolutionary completion of the embedded silencer in the chimpanzee. . Alignment of Great Apes sequences homologous to the human embedded silencer in enhancer 2 sequence. Clustal Omega, Multiple Sequence Alignment was used for alignment.

```
#  Percent Identity  Matrix - created by Clustal2.1

     1: orangutan          100.00   96.17   96.38   96.51   96.39   94.92
     2: gorilla             96.17  100.00   97.83   97.58   97.29   95.00
     3: bonobo              96.38   97.83  100.00  100.00   99.36  100.00
     4: chimpanzee          96.51   97.58  100.00  100.00   99.35  100.00
     5: human.enhancer2     96.39   97.29   99.36   99.35  100.00  100.00
     6: embedded.silencer   94.92   95.00  100.00  100.00  100.00  100.00
```

```
orangutan           agagcccctagacggcccagaggtgtctgtgttgggggacctaggtcctgaagccacctc   9065
gorilla             agagcccctatatggcccagaggtgtctgtggtgggggacctaggtcctgaagctacctc   9027
bonobo              agagcccctagatggcgcagaggtgtctgtggtgggggacctaggtcctgaagccacctc   5008
chimpanzee          agagcccctagatggcgcagaggtgtctgtggtgggggacctaggtcctgaagccacctc   149
human.enhancer2     agagcccctagatggcgcagaggtgtctgtggtgggggacctaggtcctgaagccacctc   149
embedded.silencer   -------ctagatggcgcagaggtgtctgtggtgggggacctaggtcctgaagccacctc   53
                           *** * *** ************** ********************** *****

orangutan           a-ccagaagctttcccccctgcccatccccaggtttctgggaacagattccctagggaggt   9124
gorilla             acccagaggctttcccccctgcccatccccaggtttctgggaacggattccctagggaggt   9087
bonobo              acccagaggctttcccccctgcccatccccaggtttctgggaacggattccctagggaggt   5068
chimpanzee          acccagaggctttcccccctgcccatccccaggtttctgggaacggattccctagggaggt   209
human.enhancer2     acccagaggctttcccccctgcccatccccaggtttctgggaacggattccctagggaggt   209
embedded.silencer   acccaga-----------------------------------------------------   60
                    * *****
```

Fig. S6. Alignment of the sequences of the chimpanzee and bonobo homologous to enhancer 3 with that of *SMIM45* enhancer 3. Alignment shows twelve bp mutations that distinguish the human sequence from that of the chimpanzee and bonobo. Alignment using Clustal Omega, Multiple Sequence Alignment was used for alignment.

```
#
#  Percent Identity  Matrix - created by Clustal2.1
#
#

     1: bonobo                 100.00   99.57   99.04   97.85
     2: chimpanzee              99.57  100.00   98.79   97.46
     3: human                   99.04   98.79  100.00  100.00
     4: enhancer3.NANOG.hESC    97.85   97.46  100.00  100.00


bonobo                  ggtgcaaaatgctttttaattttaatttt-tttttttttgagacagagtttcactcttgt    9107
chimpanzee              ggtgcaaaatgctttttaattttaattttttttttttttgagacagagtttcactcttgt    3094
human                   ggtgcaaaatgcttttttattttaatttt-tttttttttgagacagagtttcactcttgt    8270
enhancer3.NANOG.hESC    ----------------------------------------------agtttcactcttgt    14
                                                                      **************

bonobo                  tgcccaggctggagtgcagtggcacagtcgtggctcactggaactccacctcctgggttc    9167
chimpanzee              tgcccaggctggagtgcagtggcacagtcgtggctcactggaactccacctcctgggttc    3154
human                   tgcccaggctggagtgcagtggcacagtcgtggctcactggaactccacctcctgggttc    8330
enhancer3.NANOG.hESC    tgcccaggctggagtgcagtggcacagtcgtggctcactggaactccacctcctgggttc    74
                        ************************************************************

bonobo                  aagcaattctcttgcctcagcctgccgagtagttgggatcacaagcatgtgccaccacac    9227
chimpanzee              aagcaattctcttgcctcagcctgccgagtagttgggatcacaagcatgtgccaccacac    3214
human                   aagcaactctcctgcctcagcctgccgagtagttgggattacaagcatgtgccaccacac    8390
enhancer3.NANOG.hESC    aagcaactctcctgcctcagcctgccgagtagttgggattacaagcatgtgccaccacac    134
                        ****** **** *************************** ********************

bonobo                  ctggctaattttgtacttttagtagagacagggtttcaccatgttggtcaggctggtctt    9287
chimpanzee              ctggctaattttgtacttttagtagagacagggtttcaccatgttggtcaggctggtctt    3274
human                   ctggctaattttgtacttttagtagagacagggtttcaccatgttggtcaggctggtctt    8450
enhancer3.NANOG.hESC    ctggctaattttgtacttttagtagagacagggtttcaccatgttggtcaggctggtctt    194
                        ************************************************************

bonobo                  gaactcctgacctcaggtgacccatcctccttggcctcccaaagtgctgggattataggc    9347
chimpanzee              gaactcctgacctcaggtgacccatcctccttggcctcccaaagtgctgggattataggc    3334
human                   gaattcctgacctcaggtgacccatcctccttggcctcccaaagtgctgggatcataggc    8510
enhancer3.NANOG.hESC    gaattcctgacctcaggtgacccatcctccttggcctcccaaagtgctgggatcataggc    254
                        *** ************************************************* ******

bonobo                  atgagccattggcccggtcgcaaaatgctctttaggcattgtcttgttaaaaatgcaaag    9407
chimpanzee              atgagccattggcccggtcgcaaaatgctctttaggcattgtcttgttaaaaatgcaaag    3394
human                   atgagccattggcctggttgcaaaatgctctttaggcattgtcttgttaaactgcaaag    8570
enhancer3.NANOG.hESC    atgagccattggcctggttgcaaaatgctctttaggcattgtcttgttaaactgcaaag    314
                        ************** *** ********************************* *******

bonobo                  tacccaggctgcgtgcggtggctcacgcctgtaatcccagcactttgggaggccgaggcg    9467
chimpanzee              tacccaggctgcgtgcggtggctcacgcctgtaatcccagcactttgggaggccgaggcg    3454
human                   tacccaggctgcatgcggtggctcacgcctgtaatcccagcactttgggaggccgaggtg    8630
enhancer3.NANOG.hESC    tacccaggctgcatgcggtggctcacgcctgtaatcccagcactttgggaggccgaggtg    374
                        ************ ********************************************* *

bonobo                  ggcggatcacgaggtcaggagatcaagaccatcctggctaacacggtgaaaccccgtctc    9527
chimpanzee              ggcggatgacgaggtcaggagatcaagaccatcctggctaacacggtgaaaccccgtctc    3514
human                   ggcggatcacgaggtcaggagatcaagaccatcctggctaacacggtgaaaccccgtctc    8690
enhancer3.NANOG.hESC    ggcggatcacgaggtcaggagatcaagaccatcctggctaacacggtgaaaccccgtctc    434
                        ******* ****************************************************

bonobo                  tactaaaaatacaaaaaattagctgggcgcagtggcggtcacctgcagtcccagctactc    9587
chimpanzee              tactaaaaatacaaaaaattagctaggcgcagtggcggtcacctgcagtcccagctactc    3574
human                   tactaaaaatacaaaaaattagctgggtgcagtggcggtcacctgcagtcccagctactc    8750
enhancer3.NANOG.hESC    tactaaaaatacaaaaaattagctgggtgcagtggcggtcacctgcagtcccagctactc    494
                        *********************** ** *********************************

bonobo                  aggaggctgaggcaggagaatggtgtgaacctgggaggcggagcttgcagtgagccgaga    9647
chimpanzee              aggaggctgaggcaggagaatggtgtgaacctgggaggcggagcttgcagtgagccgaga    3634
human                   aggaggctgaggcaggagaatggtgtgaacctgggaggcggagcttgcagtgagccgaga    8810
enhancer3.NANOG.hESC    aggaggctgaggcaggag------------------------------------------    512
                        ******************
```
